# Genetic structure of Photosystem II functionality in rice unraveled by GWAS analysis

**DOI:** 10.1101/2020.09.28.317479

**Authors:** Juan Manuel Vilas, Estanislao Burgos, Maria Lucrecia Puig, Jose Colazo, Alberto Livore, Oscar Adolfo Ruiz, Fernando Carrari, Andrés Alberto Rodriguez, Santiago Javier Maiale

## Abstract

Rice production is a particularly important crop for the half-world population. Therefore, knowledge about which genes are implicated in the functionality of the Photosystem II, that are still poorly explored could collaborate in the assisted selection of rice improving. In the present study, we applied Genome wide Association Studies of PSII chlorophyll fluorescence under two contrasting environmental conditions in 283 rice accessions highly diverse. A total of 110 significant association SNP-phenotype were observed, and 69 quantitative trait loci identified with a total of 157 genes, of which 38 were highly significant, mapped spread out through rice genome. These underlying regions are enriched in genes related to biotic and abiotic stresses, transcription factors, Calvin cycle, senescence, and grain characters. The correlations analyses PSII chlorophyll fluorescence parameters and some panicle characteristics found here suggest the possibility of developing molecular markers to assist the breeding programs that improve photosynthesis and yield in rice.

**Highlight:** The genetic structure of the Photosystem II functionality in rice was studied by using genome-wide association through chlorophyll fluorescence.

## 1. Introduction

Asian rice (*Oryza sativa* L.) is one of the most important crops in the world and constitutes, approximately, the main food of the three billion humans in countries of Asia, Africa, and Latin America (Sasaki, 2005, FAO 2009).

This species is classified into five subpopulations namely: Indica, Aus, Aromatic, Tropical Japonica, and Temperate Japonica (Garris *et al*., 2005).

Rice breeding in the past has focussed on augmented the harvested index and solar radiation capture, achieving a significant increase of yield across time.

This advanced was supported, mainly, to the modified ideotype in the green revolution and improvement in agronomic practices.

Rice breeders have developed a new plant type, that improvement the past ideotype, adapting it for direct seeding. This new plant type design is characterized by reduced tillering capacity, large panicle size, lodging resistance, and specific arrangement of the three top leaves (Peng *et al*., 2004). These traits, large panicle size and arrangement of the top three leaves they aim to increment the sink size and the source of photoassimilates.

Photoassimilates are generated in the process namely photosynthesis which consists of the one light phase and another dark phase. In the light phase, photons are trapped by pigments and transport across the photosystem II (PSII) and I (PSI) in the called electron transport chain to produce NADPH and ATP.

Rice is a C3 plant where photosynthesis is limited to use solar radiation because of the importance of the photorespiration that enhancer the dissipation excess energy (Murchie *et al*., 2015; Zhu et al., 2010). Then, the control of the electron transport rate of the chain electron transport is a fundamental concept in the regulation of photosynthesis (Foyer *et al*., 2002).

Chlorophyll fluorescence is an important tool to investigate the behavior of the photosynthetic capacity and potential, this technique is as a window inside to the heart of the photosynthetic process (Stirbet and Govindjee 2011).

Different approaches have been realized to measure chlorophyll fluorescence, based on the Kautsky effect. Among them is the called OJIP analysis, which is based on the fluorescence transient of the PSII chlorophyll fluorescence and the terms concern to basal fluorescence level (F_o_), two intermediate steps namely J and I and a maximum fluorescence level (Fp = Fm) (Strasser *et al*., 2000).

Basal fluorescence level F_o_ indicated that all reaction center (RC) is open and it is assumed that at Fm levels all RC is closed.

The J step occurs at 2ms from F_o_ and is related to reduction of the primary quinone electron acceptor Q_A_ reduction, while I step occurs at the 20ms and is related to the Q_b_ reduction. Data fluorescence is recorder as F_J_ and F_I_, whilst data as fluorescence at 300μs and is used to calculate the initial slope Mo that indicates the net ratio of RCs closure.

On another hand, the difference between F_o_ and Fm is called fluorescence variable (F_v_) and the ratio between F_v_ and Fm (F_v_/F_m_) is related to the maximum quantum yield of primary PSII photochemistry, also expressed as the ratio JTR/JABS (Strasser *et al*., 2000).

Other important parameters obtained from the OJIP analysis are the area over the curve namely S_m_ that indicates a bulk of electron acceptors presented and the normalized S_m_/F_v_ indicates the energy need to close all RCs or the electron acceptors per active electron transport chain (Stirbet and Govindjee 2011).

Other authors introduced a new parameter called performance index on energy absorbed (PI_abs_). This index is calculated with three components that included density of reaction center per energy absorbed basis, energy trapping efficiency and the electron transport efficiency and can be defined as a performance index for energy conservation from photons absorbed by PSII antenna, to the reduction of Q_b_ (Srivastava *et al*., 1999).

These and other parameters calculated of the OJIP analysis allow a deep examination of the functionality of PSII and are an approach to understand the genetic variability intra/interspecies.

The genetic resources and chance to use this diversity depend on the ability to understanding the genetic base of those traits.

Quantitative trait loci (QTL) were used extensively to explore the genetic basis of the important agronomic trait. This QTL analysis developed through linkage mapping is a very time-consuming approach due to the inbreeding line must be obtained and constructing by linkage mapping. Moreover, the QTL acquired they come off the narrow genetic basis, hindering the use in the breeding program.

In recent years, the new technique to obtained QTL and molecular markers for use in plant breeding has been developed. This technique is based on the linkage disequilibrium and required a high number of genotypes with a spread genetic basis (Yang *et al*., 2018) and is called the genome-wide association study (GWAS).

In rice, numerous populations were sequenced and used in studies of GWAS as the RDP1 and RDP 1+2 consisting of 421 y 1571 accessions respectively (McCouch *et al*., 2016), two populations development in the National Center of Plant Gene Research with 533 and 950 accessions (Huang *et al*., 2012) and the 3243 rice accessions belonging to 3000 rice genome project (Li *et al*.,2014).

The study of these sequenced populations allowed the development of the great number of GWAS that covering diverse agronomic traits as heading date, plant height, grain length, spikelets for panicle, etc. (Xie *et al*., 2015; Crowell *et al*., 2016; Spindel *et al*., 2015; Begum *et al*., 2015; Zhao *et al*., 2011; Biscarini *et al*., 2016; Hu *et al*., 2016; Han *et al*., 2016), diverse biotic and abiotic stress (Li *et al*., 2017; Shakiba *et al*., 2017; Kang *et al*., 2016; Kumar *et al*., 2015; Lafarge *et al*., 2017; Shi *et al*., 2017; Kadam *et al*., 2018; Pantalião *et al*., 2016; Ma *et al*., 2016) and diverse physiological traits (Yang *et al*., 2018; Yang *et al*., 2018; Mogga *et al*., 2018; Quero *et al*., 2018).

Furthermore, currently, a great interest in describing rice phenotypes that may have better efficiency in the electrons transport (Meacham *et al*., 2017). In line with the above, there are reports in rice that describe the relationship between PSII activity and grain yield (Wang *et al*., 2014; Zhang *et al*., 2015).

In the present study, we applied GWAS mapping to investigate the genetic architecture of PSII efficiency under two environmental conditions and evaluated the interaction between the behaviour of PSII and yield components. We analyzed PSII efficiency in RDP 1 population with highly diverse rice genotypes; these populations were genotyped with 700,000 SNPs markers (Crowell *et al*., 2014). Indeed, these results are discussed in the context of the rice PSII efficiency and they translated into yield.

## 2. Material and methods

### 2.1 GWAS germplasm selection

A collection of 283 accessions, belonging to the RDP-1 panel, representing the five major subpopulations in *O. Sativa* genotyped to 700,000 SNPs (Crowell *et al*., 2016) was used. The accessions selected from the rice subpopulations were 45 *aus*, 59 *indica, 58 temperate japonica, 76 tropical japonica, 8 aromatics, and 37 admixed*.

### 2.2. Experimental design and locations

The seeds were sown in Petri dishes on filter paper, rinsed with 7 mL carbendazim 0.025 %p/v, and incubated at 30°C in darkness during three days until germination. The seedlings were cultured in a growth chamber with 28/24 °C day/night temperatures, 12-12h photoperiod 240 μmol photons m^−2^ s^−1^ of photosynthetically active radiation (PAR) and, 60% average humidity. The seedling was growing in hydroponically conditions with nutritive Yoshida solutions (Yoshida *et al*., 1976). After two weeks germination seedling was transplanting to field conditions in pots containing mineral soil. The accessions were planted in a randomized complete form with three replications.

Field experiments were conducted at two locations, INTECH, Chascomús-Buenos Aires (35.623048 −57.994167, CH), and EEA INTA Concepción del Uruguay, Entre Ríos (32.490254, −58.348855, ER). Both experiments were performed in the summer season 2017/2018. The physiological measurements were performed in plants of 11 old-weeks and all plants are in the tillering stage.

### 2.3 Chlorophyll transient fluorescence measurements (OJIP test) and SPAD

Fluorescence transient was measured in the uppermost fully expanded leaf. Measurements were taken with a plant efficiency analyzer (Handy PEA fluorometer, Hansatech Instruments Ltd., King’s Lynn, Norfolk, UK), according to the protocol used for the so-called double-hit experiments (Appenroth *et al*., 2001). Leaves were dark-adapted by 20 minutes using a leaf clip provided by the manufacturer. Then, leaves were exposed twice for 1 s with 3000 μmol photons m^−2^ s^−1^ (650 nm peak wavelengths) with a dark interval of 0.5 seconds. In this line, the data become for the first hit was extracted and used as a phenotypic fluorescence trait.

The fraction of Q_b_ reducing centers (Q_b_RC) can be calculated as follows:

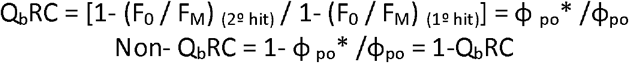

The PEA plus software (PEA Plus software, Hansatech Instruments Ltd., UK) was used to analyze the PSII parameters according to Strasser *et al*. (2000). The different OJIP parameters are present in Table 1.

**Table 1:**
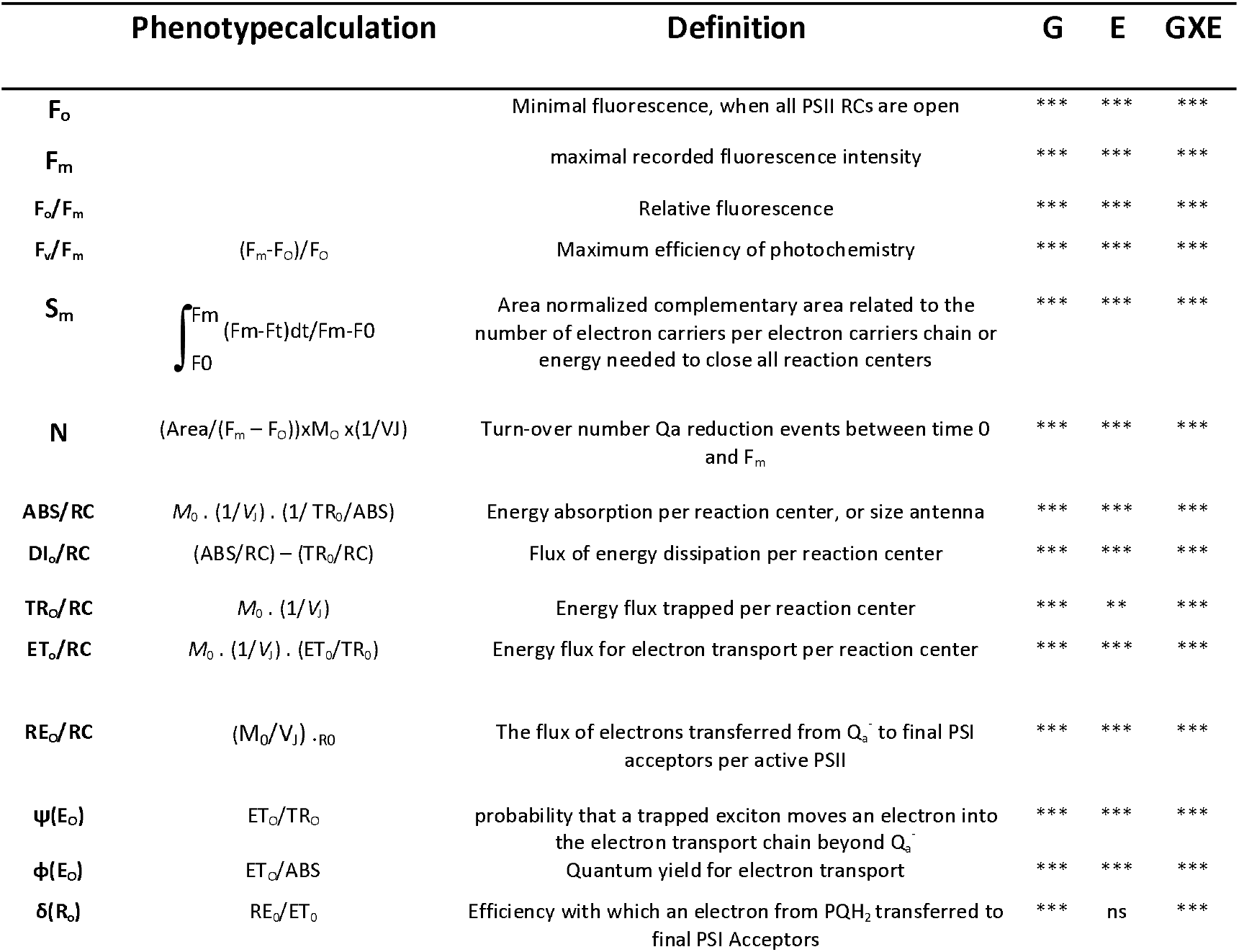
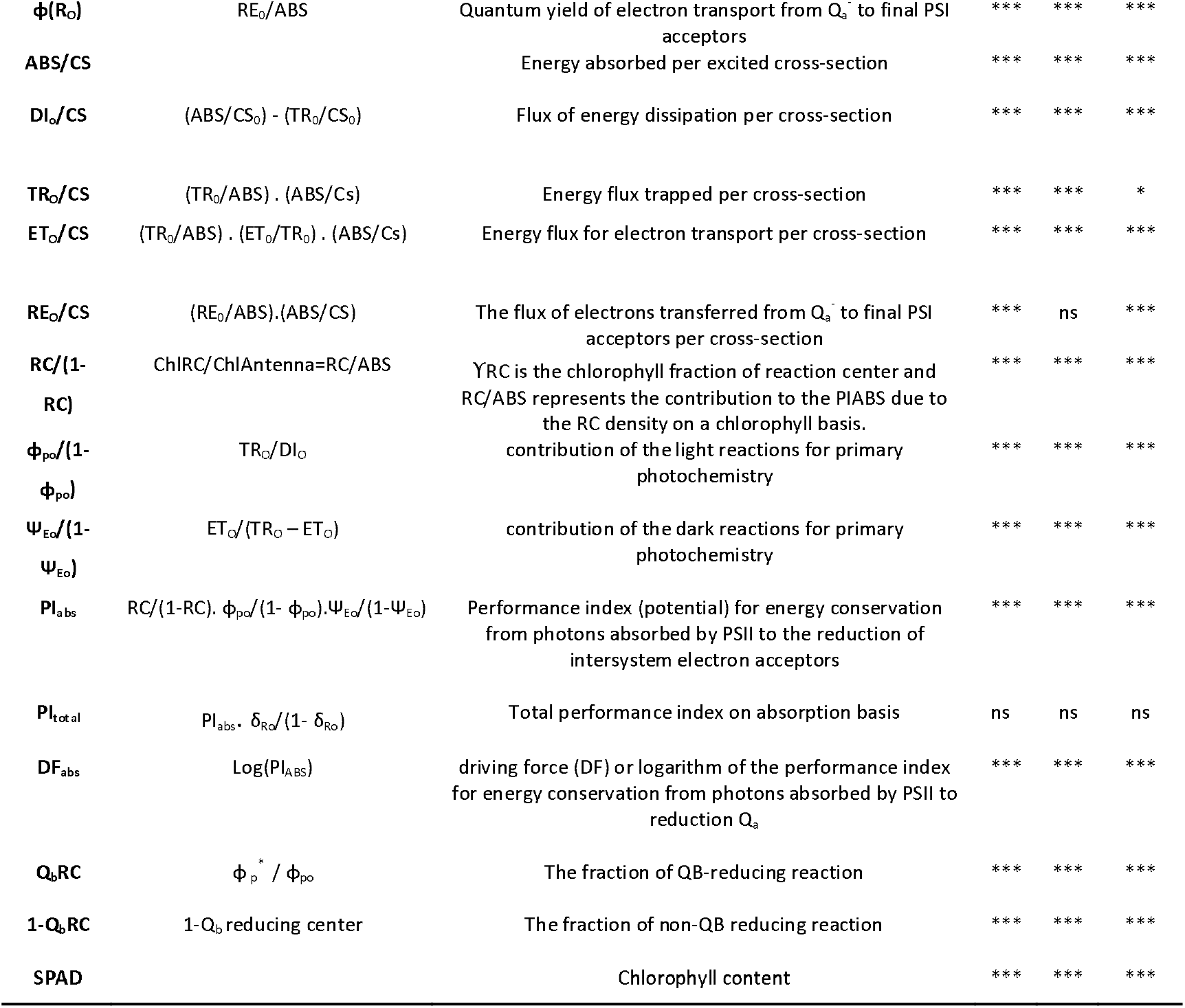
Calculations and definitions of fluorescence phenotypes (Strasser *et al*. 2000) and ANOVA analyze of environment (E), genotype (G) and genotype by environment effects on fluorescencephenotypes. Significance of the P-values is given as: *** p< 0.001, **p< 0.01, *p< 0.05, ns (no significance).

To estimate leaf total chlorophyll content, a SPAD chlorophyll meter was used (Cava-devices, CABA, AR). For each leaf, the chlorophyll content was estimated as the mean of five chlorophyll content measurements at different positions in the middle section of the leaf.

### 2.4 Determination of yield components

Plants were harvested at the maturation stage. The panicles were threshed manually and processed by a hulling machine JLGJ-45 (JinGao, Cn) and the following yield components were measured: the weight of one thousand grain (WTG), number of panicle per plant (PN), number of spikelets per panicle (NSP), panicle weight (PW) calculated as the weight of filled grains per panicle, percentage of fertile and infertile spikelets (%F and %In respectively). All dates are an average of total panicles per plant replications.

### 2.5 GWAS mapping

The GWA studies were performed based on the HDRA data set consisting of 700,000 SNPs (McCouch *et al*., 2016). Phenotypic grand means for each variety were used in the mapping models. GWAS was performed using a linear mixed model with EMMAX algorithm (Kang *et al*., 2010), which takes the underlying population structure into account by including two kinship matrix (hBN and hIBS) as a covariate.

The maximum rates of missing data were fixed at 10% per accessions and 25% per SNP. A minor allele frequency threshold of 0.05 (MAF<0.05) was applied to discard markers with very rare alleles according to Aulchenko *et al*. (2007). The significance threshold was set at p< 1.28 x 10^−7^ for every trait, using Bonferroni criteria.

### 2.6 Linkage disequilibrium

GWAS Linkage disequilibrium (LD) between markers on each chromosome was calculated using pairwise r2 between SNPs. First, we performed LD, calculated using the –r2 –Id-window 99999 – Id-window-r2 0 commands in PLINK (version 1.9) (Chang *et al*., 2015).

The critical threshold used was 0.2 (McCouch *et al*., 2016). Then, for each significant marker, we computed LD upstream and downstream on the same chromosome as a mean LD value through plotting r2 1,000 SNPs of each chromosome versus genetic distance between markers we defined both boundaries of the confident interval as the intersection of the ‘critical LD’ threshold and the fitted curve of r2 regression (See Supplementary. Fig.1). A nonlinear model described by Remington *et al*. (2001) was used to fit the LD decay. The R function NLS (nonlinear least-squares method) was used to fit the model. The number of genes within each interval was identified from the rice genome.

### 2.7 Statistical analyses

Histograms, boxplots, correlations and GWAS analyses were constructed using phenotypic grand means for each variety. Manhattan plot and Qqplots (Suppl. Figure 2 and 3) were created using the package qqman() in R (Rebolledo *et al*., 2015). P-values for Pearson’s correlation coefficients were calculated with a two-sided t-test using the cor.test() function in R.

Analysis of variance was performed for genotype, environment, and genotype x environment using the lme4 function in R (Bates *et al*., 2015).

### 2.8 Gen candidate analysis

A list of the candidate gene was constructed in the base to gen functional notation utilizing the Rice Gene Annotation Project database (http://rice.plantbiology.msu.edu/cgi-bin/batch_download.pl).

Also, an enrichment analysis of the GO terms (Gene Ontology) was performed, using the agriGO analysis tool (http://systemsbiology.cau.edu.cn/agriGOv2; Du *et al*., 2010). In this analysis, the list of gen candidates derived from the GWAS mapping was used utilizing the Oryza Sativa LMSU database with agriGO as a reference. The analysis was carried out using Fisher’s statistical method to detect significantly enriched ontological terms. The transcription factors (FT) identified were classified according to the PlnTFDB web tool (the PlantTranscription Factor Database; http://plntfdb.bio.uni-potsdam.de/v3.0; Pérez-Rodríguez *et al*., 2009)

## 3. Results

### 3.1 Phenotype and population structure

The rice diversity panel RDP-1 consists of 435 inbred accessions of *O. sativa* that come from 82 different countries (Zhao *et al*., 2011). In this study, a part of the RDP-1 was cultivated under field conditions in two different environmental locations, Chascomús (CH) and Entre Ríos (ER). Then, 283 plant accessions (from those above), were phenotyping at vegetative stage for photosynthetic traits (PT) by measuring the fluorescence transient of chlorophyll a and the SPAD index. On the other hand, a principal component analysis (PCA) was used to describe the overall genetic variation in the RDP-1. The population presented a clear structuring of the accessions analyzed caused by the environmental condition (Fig. 1). The top two principal components (PCs) explain 95.17% of the genetic variation within the panel (Figure 1). A total of 29 PT were measured for 283 accessions in RDP-1 at the vegetative stage and they were used for the GWAS analysis, in which all rice subpopulations were represented: 45 aus, 59 indica, 58 temperate japonica, 76 tropical japonica, 8 aromatic and 37 admixed. A brief description of the traits evaluated is summarized in Table 1. In this sense, the analysis of variance (ANOVA) showed significant effects on the genetic component (G), the environmental factor (E), and their interaction (GxE) (Table 1). However, some exceptions were the RE_o_/CS and δ(R_o_) traits, that not showed significant E effects. In line with this the PItotal not presented significant differences in G, E and GxE interaction (Table 1).

**Figure 1:**
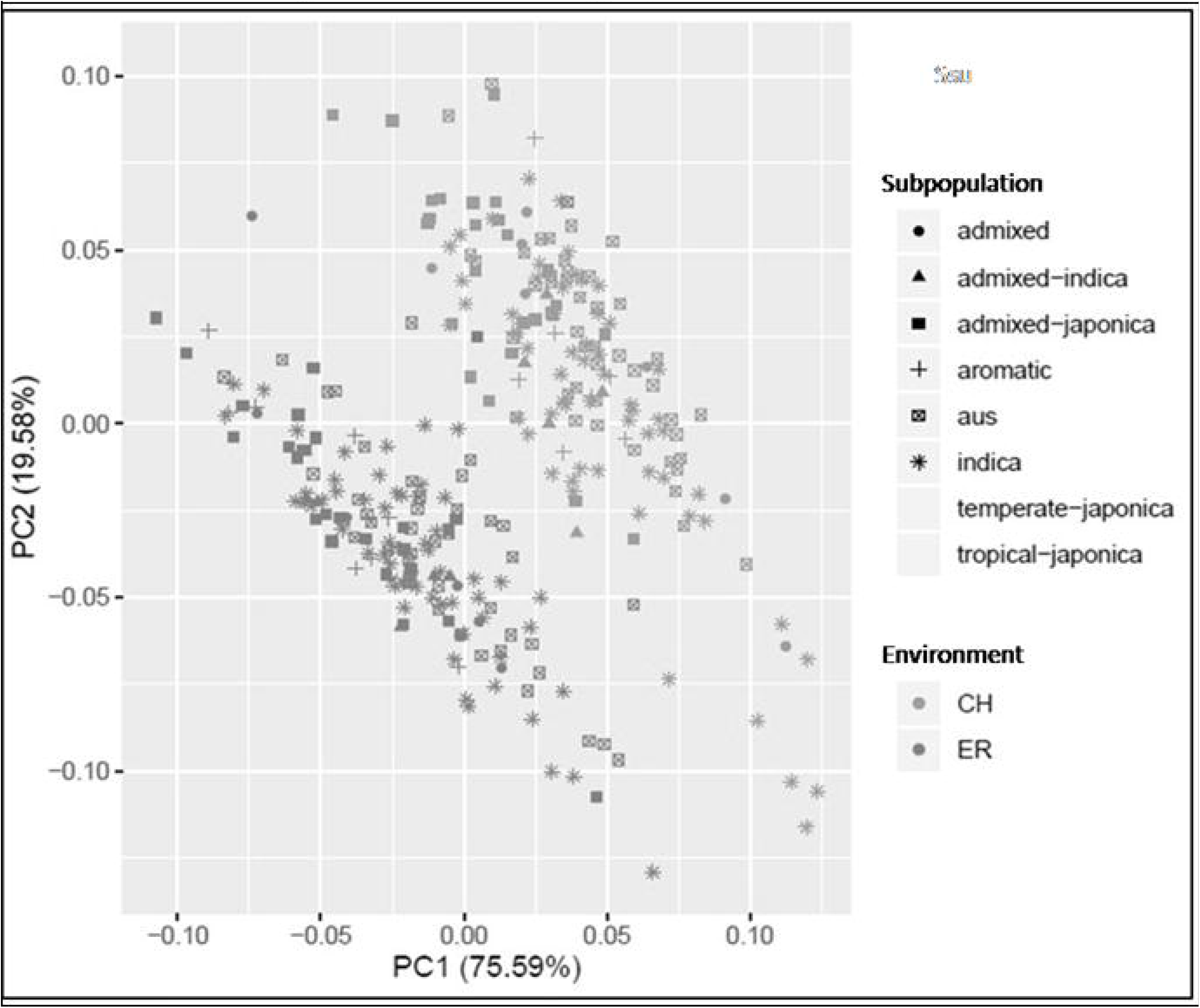
Genetic structure of the population: PCA analysis for the first two PCs, the CH environment is represented in light gray and the ER environment is represented in dark gray.

### 3.2 Genome-wide association analysis

After filtering, the set of markers was constituted of 388,084 SNPs with MAF ≥ 5%, within the 12 rice chromosomes. A total of 110 significant SNPs-trait associations were identified, intersecting both sets of significant associations (obtained with hIBS and hBN matrices, mention above). Moreover, 11 of these significant SNP-trait associations corresponded to the cultivation plants in CH environment, while 99 significant SNP-trait associations belonged to plants grown in the ER environment (Table 2). Interestingly these associations being unique for each environment, presenting none SNP in common between CH and ER. Likewise, of the total fluorescent phenotypes evaluated here, 5 of them registered significant associations in both environments with SNPs (F_o_/F_m_, F_v_/F_m_, DI_o_/RC, δ(R_o_), and PI_abs_ (Table 2) in both environments of the PT analyzed, only the chlorophyll fluorescence phenotypes showed significant associations with SNPs. In line with this, 6 significant fluorescent phenotype associations were detected in CH environment, while in ER environment, a total of 13 phenotypes were significantly associated with some SNPs (Fig. 2A-B; Table 2). The plants growing in the CH environment showed a lower mean value of the F_v_/F_m_ and PI_abs_ traits than ER environment, with differences of 7.23% and 57.27% respectively (Fig. 2). However, other phenotypes like F_o_/F_m_, DI_o_/RC, and δ(R_o_) of the plants grown in CH showed higher average values compared to the ER environment. These differences being 26.17% for the F_o_/F_m_, 31.55% for the DI_o_/RC, and 31.77% for the δ(R_o_) (Fig. 2).

**Figure 2:**
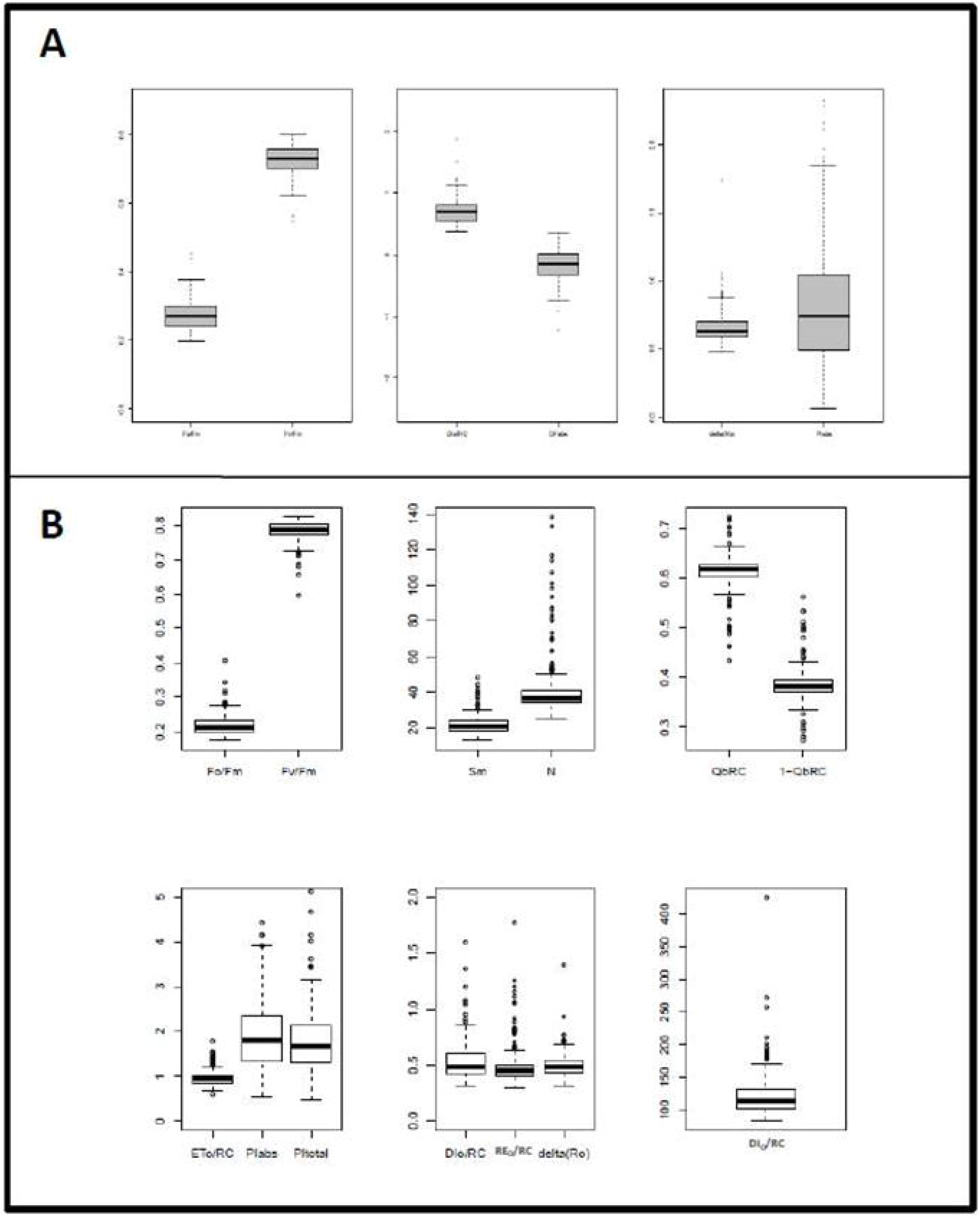
Box plots of fluorescence phenotypes with significant associations in GWAS analysis. The line inside the box represents the mean value of each phenotype, the top and bottom of each represent the 1st and 3rd quartiles. Open circles represent outliers within the dataset. (A). Gray boxes phenotypes corresponding to the environment CH. (B). White boxes phenotypes corresponding to the ER environment.

**Table 2:**
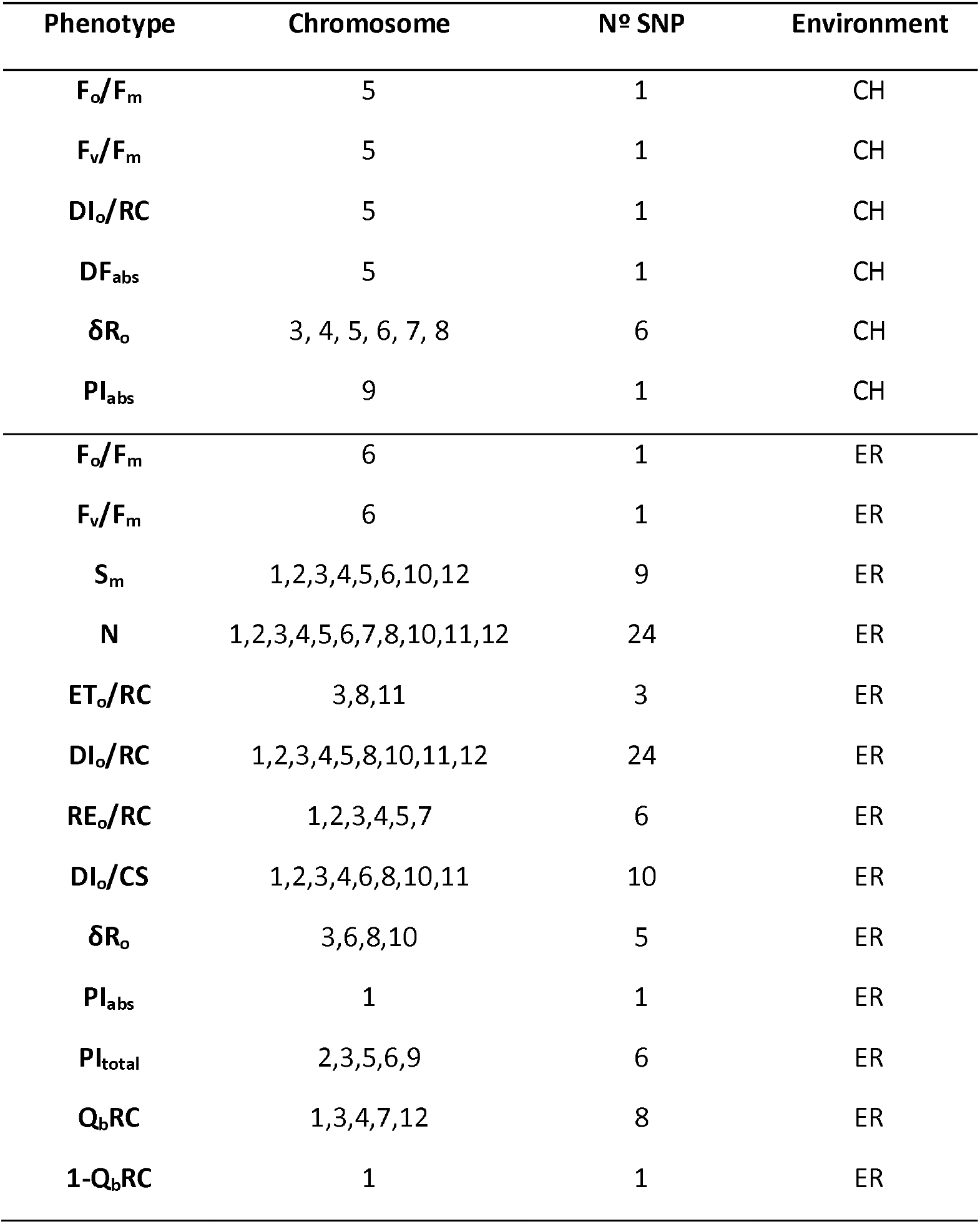
Summary of significative associations between SNP and Chlorophyll Fluorescence phenotypes in the environment CH and ER.

| Phenotype | Chromosome | Nº SNP | Environment |
| --- | --- | --- | --- |
| $F_o/F_m$ | 5 | 1 | CH |
| $F_v/F_m$ | 5 | 1 | CH |
| $DI_o/RC$ | 5 | 1 | CH |
| $DF_{abs}$ | 5 | 1 | CH |
| $\delta R_o$ | 3, 4, 5, 6, 7, 8 | 6 | CH |
| $PI_{abs}$ | 9 | 1 | CH |
| $F_o/F_m$ | 6 | 1 | ER |
| $F_v/F_m$ | 6 | 1 | ER |
| $S_m$ | 1,2,3,4,5,6,10,12 | 9 | ER |
| $N$ | 1,2,3,4,5,6,7,8,10,11,12 | 24 | ER |
| $ET_o/RC$ | 3,8,11 | 3 | ER |
| $DI_o/RC$ | 1,2,3,4,5,8,10,11,12 | 24 | ER |
| $RE_o/RC$ | 1,2,3,4,5,7 | 6 | ER |
| $DI_o/CS$ | 1,2,3,4,6,8,10,11 | 10 | ER |
| $\delta R_o$ | 3,6,8,10 | 5 | ER |
| $PI_{abs}$ | 1 | 1 | ER |
| $PI_{total}$ | 2,3,5,6,9 | 6 | ER |
| $Q_b/RC$ | 1,3,4,7,12 | 8 | ER |
| $1-Q_b/RC$ | 1 | 1 | ER |

On the other hand, among the 110 SNPs mapped here for both environments, 70 SNPs were associated with a single locus trait (Table 2). In this sense, others 11 associations found for CH, the F_o_/F_m_, F_v_/F_m_, DI_o_/RC, and DF_abs_ phenotypes mapped in the same SNP, SNP-5.11015630 (Supplementary Table S1), while the 6 associations found for the δ(R_o_) phenotype were unique for this parameter, as well as the only SNP identified for the PI_abs_ (Table 2).

Alternatively, in ER environment, 63 SNPs were associated with a single locus respectively (Table 2), as for the CH environment, PI_abs_ phenotype mapped in a single significant SNP, SNP-1.35479825 (Supplementary Table S1), while other 12 and 4 SNPs were associated together with two and three fluorescence traits respectively (Supplementary Table S1). Among the 28 fluorescence traits analyzed here, the turnover of Q_a_ (N) and energy dissipated by active reaction centers of FSII (DI_o_/RC), were associated with the higher number of SNPs (24 SNPs, Table 2).

Likewise, a total of 69 QTLs were identified for 14 of 29 PT analyzed in both environments tested and belonged to three classes. The first class of QTL was defined as single SNP associated with a unique phenotype, the second class of QTL were those identified SNPs associated with more than one phenotypes (multi-type QTL). Finally, the third class of QTL found corresponded to SNPs mapped in chromosome region, where two or more SNPs separated by a distance of ~ 55 kb and these were associated with a particular phenotype (QTL multi SNP). In line with this, the CH environment, presented a single multi-type QTL on chromosome 5 (Fig. 3A), where a single SNP was associated with the F_o_/F_m_, F_v_/F_m_, DI_o_/RC, and DF_abs_ traits (SNP-5.11015630).

**Figure 3:**
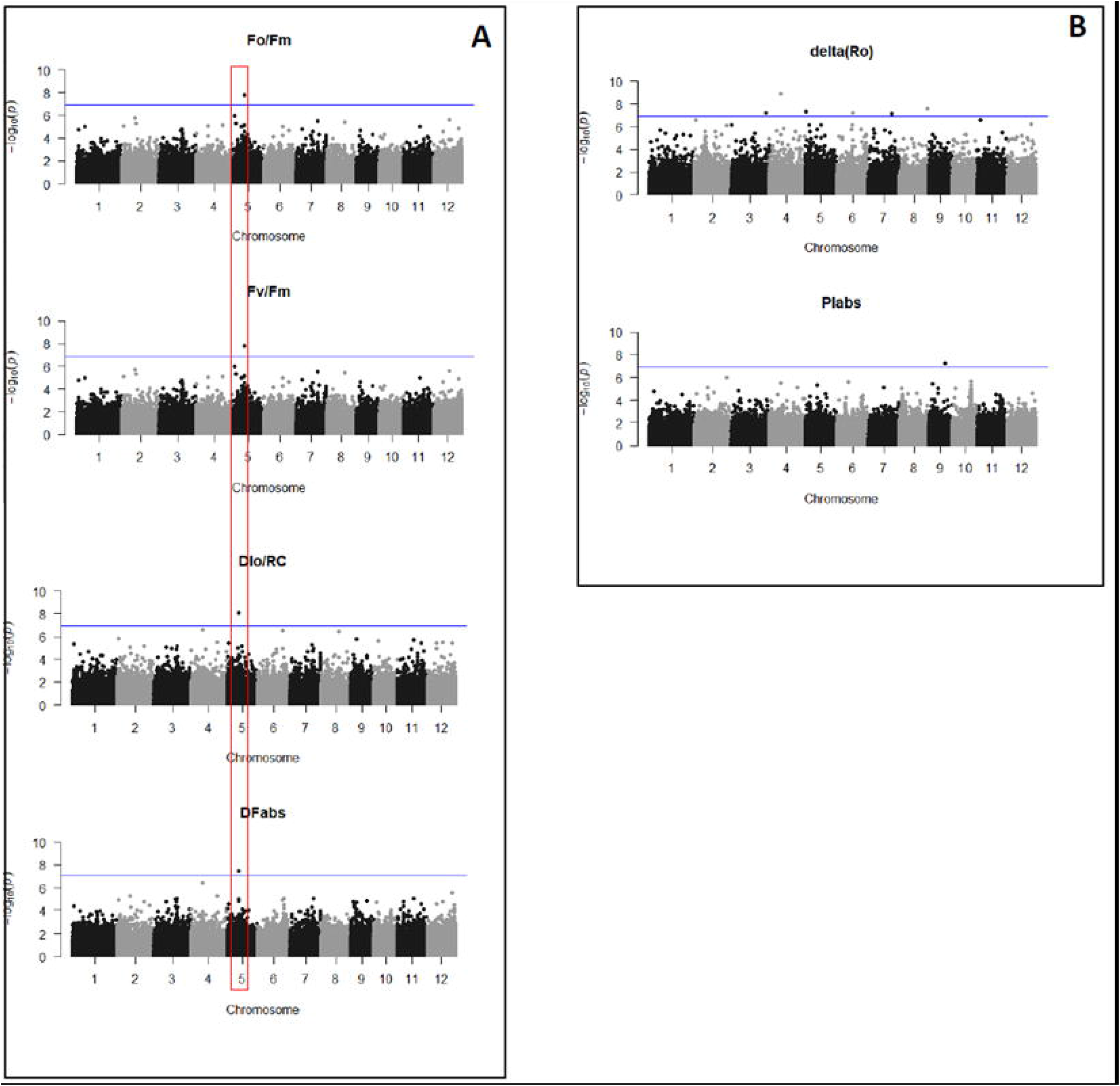
Manhattan plots for the significant phenotypes found using GWAS for the environment CH. (A). Plots for the F_o_/F_m_, F_v_/F_m_ DI_o_/RC, and DF_abs_ phenotypes. (B). Plots for the δ(R_o_) and PI_abs_ phenotypes. The box in red identifies the multi-phenotypic QTL and each SNP is represented by a gray or black dot. The x-axis shows each of the 12 rice chromosomes, the y-axis represents the negative base ten logarithms (-log10) of each p-value corresponding to a particular SNP. The blue line that cuts the y-axis represents the significance threshold value for each SNP.

On the other hand, for the ER environment, 4 multi-type QTLs were identified on chromosomes 2, 3, 4, and 6 (Fig. 4A), where the RE_o_/RC, N, and S_m_ phenotypes shared the SNP SNP-2.18755532 (Supplementary Table S1). At the same time, the δ(R_o_), DI_o_/CS, and DI_o_/RC phenotypes shared the SNP, SNP-3.7631091. In turn, the DI_o_/RC phenotypes shared SNP-4.3532129 with the N and RE_o_/RC phenotypes. Finally, the F_o_/F_m_, F_v_/ F_m_, and δ(R_o_) traits were jointly associated with the locus SNP-6.21617771 (Fig. 4A; Supplementary Table S1).

**Figure 4:**
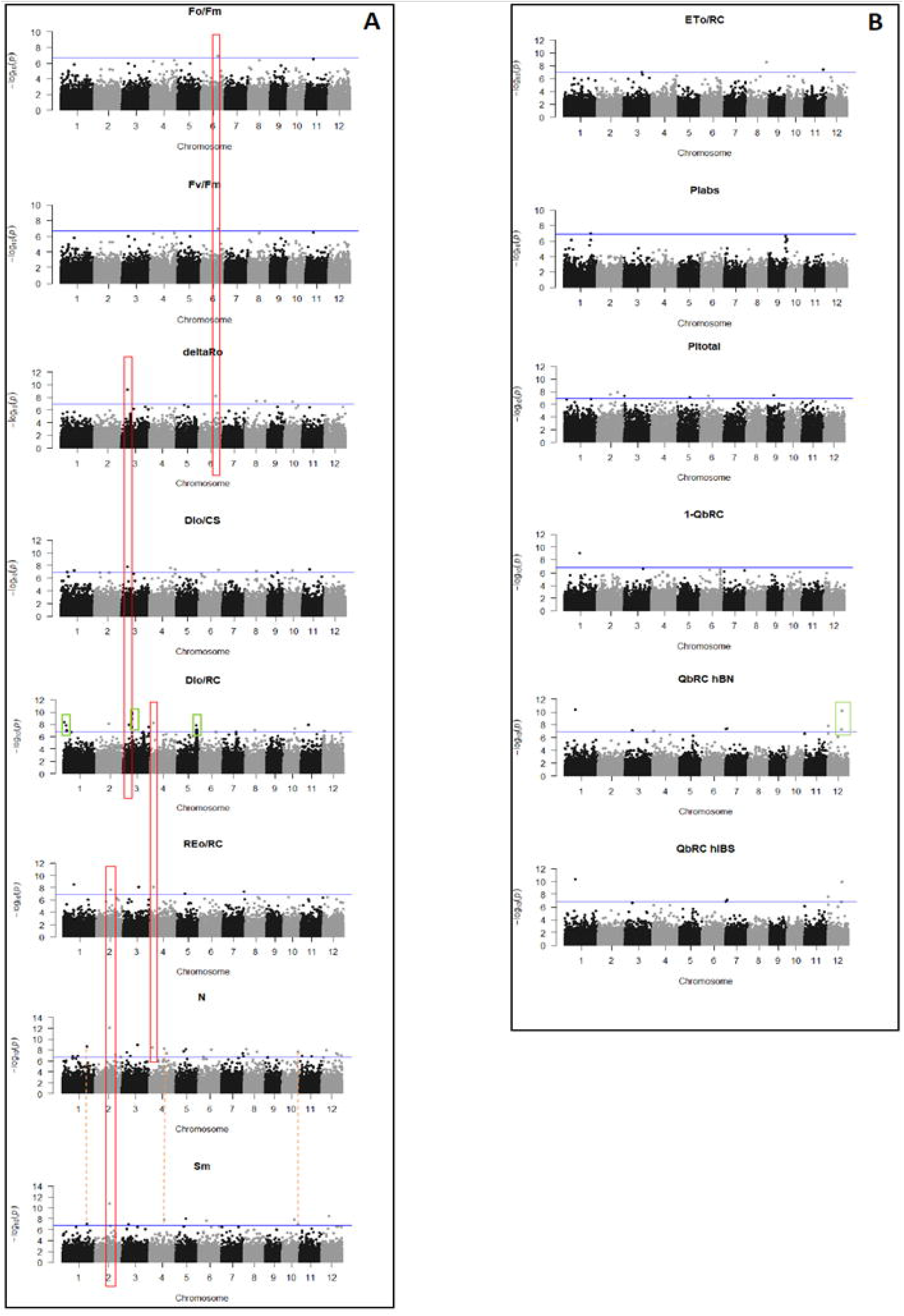
Manhattan plots for the significant phenotypes found by GWAS for the ER environment. (A). Plots for the F_o_/F_m_, F_v_/F_m_, δ(R_o_), DI_o_/RC, DI_o_/CS, RE_o_/RC, N and S_m_ phenotypes, (B). Plots for the ET_o_/RC, PI_abs_, PI_total_, Q_b_RC, and 1-Q_b_RC phenotypes (hBN and HIBS). The box in red and the orange dotted line identify multi-phenotyp QTL, green box identifies the multi SNP QTLs, and each SNP is represented by a gray or black dot. The x-axis shows each of the 12 rice chromosomes, the y-axis represents the negative base ten logarithms (-log10) of each p-value corresponding to a particular SNP. The blue line that cuts the y-axis represents the significance threshold value for each SNP.

Besides, another 4 QTL multi-types were identified for S_m_ and N on chromosomes 1, 4, 5, and 10 (SNPs 1.32181171, 4.19830956, 5.13606968 and 10.2073752). In addition one more multi-type QTL of interest was found for the energy dissipation phenotypes DI_o_/RC and DI_o_/CS, on chromosome 11; SNP11.9898109 (Fig. 4A). Finally, 5 QTL multi SNPs were detected, 3 of these were associated with DI_o_/RC trait on chromosomes 1, 3, and 5 (Fig. 4A), and other one for the Q_b_RC onto chromosomes 12 (Fig. 4B). The remaining 59 QTLs identified corresponded to the single SNP class (Fig. 4A-B).

### 3.3 Identification of candidate genes

The genome browser feature of Rice base “Rice Annotation Project” http://rice.plantbiology.msu.edu/index.shtml, was used to identify candidate genes based on the MSU7 gene annotation. The selection of candidate genes was underlined for each QTL accordingly the linkage disequilibrium (LD) (Supplementary Table S2). These genes were classified accordingly their Gene ontology (GO) term. Subsequently the enrichment analysis of the list of total candidate genes showed a significant proportion of genes involved on molecular functions related to cell biosynthesis (GO: 0044249), metabolic regulation processes (GO: 0019222), regulation of gene expression and macromolecular processes (GO: 0010468, GO: 0060255), processes related to cell death (GO: 0008219), macromolecule cell biosynthesis (GO: 0034645), and macromolecule biosynthesis (GO: 0009059) also showed significant enrichment (Table 3). Furthermore the most represented terms linked to biological processes were related to hydrolysis processes of acids, phosphate and ester groups (GO: 0016817, GO: 0017111, GO: 0016818, GO: 0016462 and GO: 0016788) and nucleotide-binding (GO: 0000166).

**Table 3:**
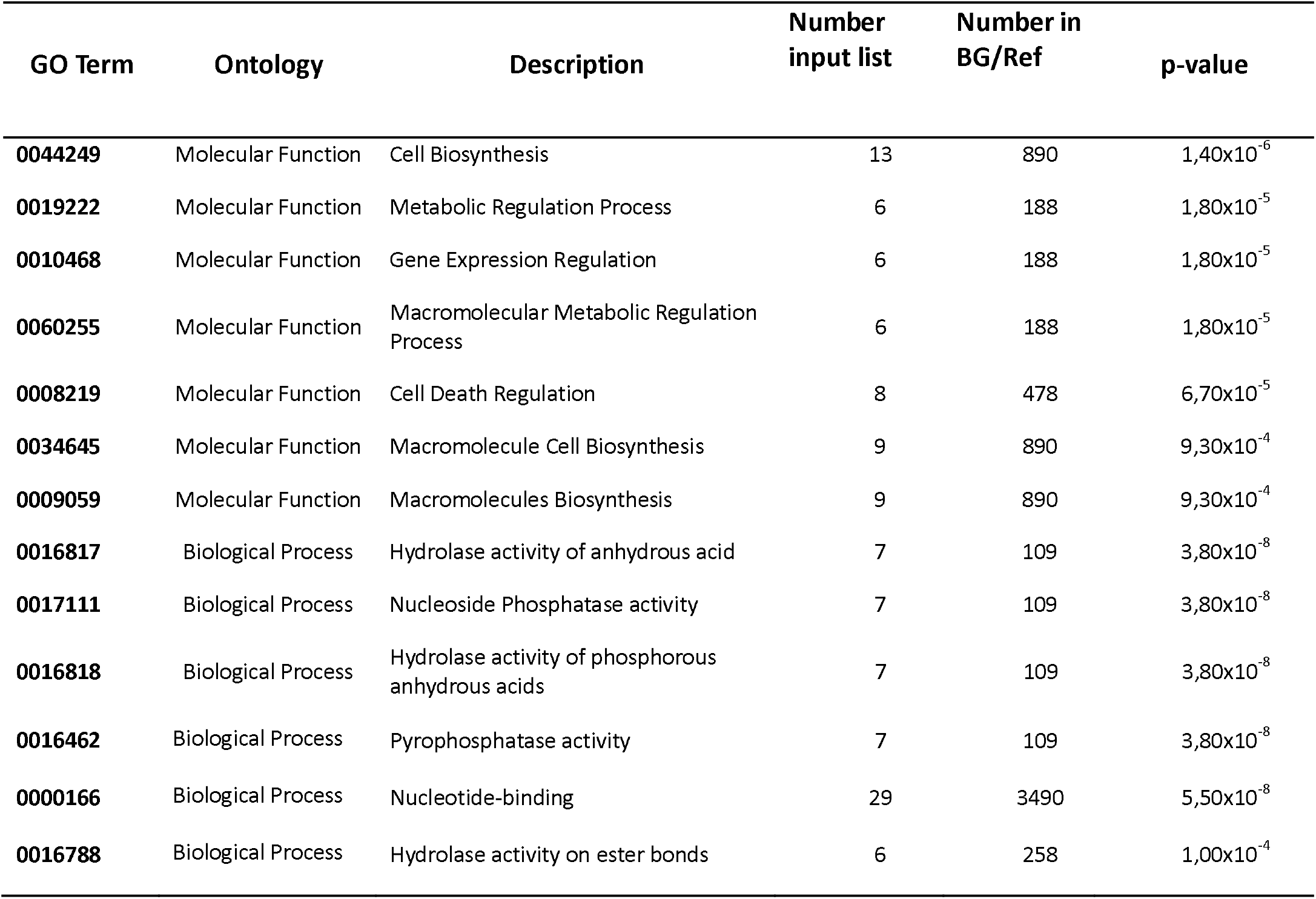
Enrichment analysis of GO terms based on identified candidate genes: The most statistically representative Go terms are shown.

Based on these most representative GO terms, 38 candidate genes were highlighted in Table 4 from the 157 total candidate genes mapped (Supplementary Table S2).

**Table 4:** Summary of the most important candidate genes identified by GWAS analysis, linked to the most representative Go Terms, Phenotype, Environment, and SNP.

| SNP | GO Term | ID | Functional annotation | Environment | Phenotype |
| --- | --- | --- | --- | --- | --- |
| SNP-9.16154636 | GO: 0000166 | LOC_Os09g26620 | Auxin repressed protein | CH | Plabs |
| SNP-5.920701 | GO: 0000166 | LOC_Os05g02670 | Kinesin motor domain protein | CH | $\delta(R_o)$ |
| SNP-10.20737522<br>SNP-3.11832167<br>SNP-3.11839954<br>SNP-3.11840203 | GO:0044249 | LOC_Os10g39030 | Homeobox domain | ER | $S_m$<br>N<br>$DI_o/RC$ |
| SNP-5.13606968 | GO:0044249 | LOC_Os05g23780 | OsMADS70 family of genes - MADS-box with expressed M-alpha type-box | ER | $S_m$<br>N |
| SNP-1.32181171 | GO:0044249 | LOC_Os01g55870 | Chloroplast precursor of Corismatemutase | ER | $S_m$<br>N |
| SNP-1.32181171 | GO:0044249 | LOC_Os01g55890 | Amino acid kinase | ER | $S_m$<br>N |
| SNP-5.26255003.<br>SNP-5.26255003 | GO:0044249 GO: 0000166 | LOC_Os05g45340<br>LOC_Os05g45340 | ATP binding protein | ER | $DI_o/RC$ |
| SNP-5.25755857 | GO:0044249 | LOC_Os05g44360 | Oligosaccharide Transferase | ER | $DI_o/RC$ |
| SNP-3.9505313 | GO:0044249 | LOC_Os03g17120 | Arginine biosynthesis, argJbifunctional protein | ER | $Q_bRC$ |
| SNP-3.9505313 | GO:0044249 | LOC_Os03g17130 | Transcription factor b Helix-loop-helix | ER | $Q_bRC$ |
| SNP-5.25752878<br>SNP-5.25755857 | GO:0044249 | LOC_Os05g44390 | Peptidyl-tRNA hydrolase | ER | $DI_o/RC$ |
| SNP-5.25752878<br>SNP-5.25755857 | GO:0044249 | LOC_Os05g44400 | GATA zincfingerdomain | ER | $DI_o/RC$ |
| SNP-3.11913691<br>SNP-3.11909250 | GO:0044249 | LOC_Os03g20970 | Phospholipid transporter of type ATPase 1 | ER | $DI_o/RC$ |
| SNP-3.11832167<br>SNP-3.11840203<br>SNP-3.11839954 | GO:0044249 | LOC_Os03g20900 | Myb transcription factor | ER | $DI_o/RC$ |
| SNP-1.672866 | GO: 0000166 | LOC_Os01g02250 | RGA-1 | ER | $DI_o/RC$ |
| SNP-1.2950882 | GO: 0000166 | LOC_Os01g06240 | Protein kinase | ER | $DI_o/RC$ |
| SNP-1.3777578 | GO: 0000166 | LOC_Os01g07870 | Transporter family protein type A | ER | $DI_o/RC$ |
| SNP-1.20577084 | GO: 0000166 | LOC_Os01g36920 | ATP-dependent DEAD-box RNA helicase | ER | N<br>$1-Q_bRC$ |
| SNP-1.32181171 | GO: 0000166 | LOC_Os01g55880 | DNA hemimethylated domain-binding protein | ER | $S_m$<br>N |
| SNP-2.18755532 | GO: 0000166 | LOC_Os02g31290 | AML1 | ER | $RE_o/RC$ $S_m$<br>N |
| SNP-3.6370379 | GO: 0000166 | LOC_Os03g12150 | Serine/threonine receptor protein kinase precursor | ER | N |

| SNP | GO Term | ID | Functional annotation | Environment | Pheno type |
| --- | --- | --- | --- | --- | --- |
| SNP-3.33035702 | GO: 0000166 | LOC_Os03g59600 | Mitochondrial RhoGTPase1 | ER | DI<sub>o</sub>/RC |
| SNP-3.33935702 | GO: 0000166 | LOC_Os03g59620 | Phospholipase of the patatin family | ER | DI<sub>o</sub>/RC |
| SNP-5.13606968 | GO: 0000166 | LOC_Os05g23800 | Motif for RNA recognition | ER | S<sub>m</sub><br>N |
| SNP-5.25752878<br>SNP-5.25755857 | GO: 0000166 | LOC_Os05g44340 | Heat shock protein 101 | ER | DI<sub>o</sub>/RC |
| SNP-5.25860213<br>SNP-5.25870238 | GO: 0000166 | LOC_Os05g44560 | Protein Kinesin motor domain | ER | DI<sub>o</sub>/RC |
| SNP-5.26037406<br>SNP-5.26067844 | GO: 0000166 | LOC_Os05g44922 | 6-phosphofructokinase | ER | DI<sub>o</sub>/RC |
| SNP-5.26037406<br>SNP-5.26067844 | GO: 0000166 | LOC_Os05g44930 | Protein kinase type receptor | ER | DI<sub>o</sub>/RC |
| SNP-5.26067844 | GO: 0000166 | LOC_Os05g44940 | Protein containing kinase domain | ER | DI<sub>o</sub>/RC |
| SNP-5.26067844 | GO: 0000166 | LOC_Os05g44970 | Senescence receptor precursor of the Serine / Threonine protein kinase type | ER | DI<sub>o</sub>/RC |
| SNP-6.17586886 | GO: 0000166 | LOC_Os06g30430 | RPM1 disease resistance protein | ER | N |
| SNP-8.17383104 | GO: 0000166 | LOC_Os08g28470 | RPM disease resistance protein | ER | N |
| SNP-11.9898109 | GO: 0000166 | LOC_Os09g02400 | RNP RNA binding region | ER | DI<sub>o</sub>/RC |
| SNP-9.8690732 | GO: 0000166 | LOC_Os09g14660 | Legume lectins containing beta domain proteins | ER | PI<sub>total</sub> |
| SNP-12.1522824 | GO: 0000166 | LOC_Os12g03750 | Leucine-rich protein family in | ER | DI<sub>o</sub>/RC |
| SNP-12.18066755 | GO: 0000166 | LOC_Os12g30150 | CAMK_CAMK_del type 47 - CAMK includes calcium / calmodulin-dependent protein kinases | ER | Q<sub>b</sub>RC |
| SNP-12.18757106<br>SNP-12.19687181<br>SNP-12.19687181 | GO: 0000166 | LOC_Os12g31200<br>LOC_Os12g32680 | Domain that contains NB-ARC | ER | Q<sub>b</sub>RC |
| SNP-12.19687181 | GO: 0000166 | LOC_Os12g32670 | CC-NBS-LRR resistance protein | ER | N |

Of those 38 candidate genes 8 genes corresponded to genes associated with response to biotic and abiotic stress (LOC_Os01g02250, LOC_Os03g59600, LOC_Os05g44340, LOC_Os06g30430, LOC_Os08g28470, LOC_Os09g26620, LOC_Os12g03750, and LOC_Os12g30150). Also, 5 genes were identified as transcription factors (LOC_Os03g17130, LOC_Os03g20900, LOC_Os05g23780, LOC_Os05g44400 and LOC_Os10g39030), one gene was assigned to the Calvin Cycle (LOC_Os05g44922). Likewise, three identified genes are linked to functions associated with grain characters (LOC_Os05g02670, LOC_Os05g44560, and LOC_Os10g39030); and finally, a gene with functional notation linked to senescence processes (LOC_Os05g44970). Based on these results, it could be said that the genetic architecture of the functionality of the PSII can be associated at the genetic level mainly with processes linked to grain filling among other processes.

### 3.4 Relationships between panicle traits and FSII parameters

Six panicle traits were evaluated: Number of Panicle per Plant (PN), Thousand Grain Weight (WTG), Fertility Percentage (%F), Calculated Infertility Percentage (%In), Number of Spikelet per Panicle (NSP) and Panicle Weight (PW).

These panicle traits showed different distribution depending on the environmental condition. Plants grown in the environmental conditions of ER showed higher NP, NSP, %In, and PW compared with plants grown in the CH; these differences were 15.1%, 45.6%, 26.1%, and 27.4% respectively (Fig. 5). Meanwhile, the WTG and %F traits were 10.3% and 19.80% lower for the ER environment compared to the CH environment (Figure 5).

**Figure 5:**
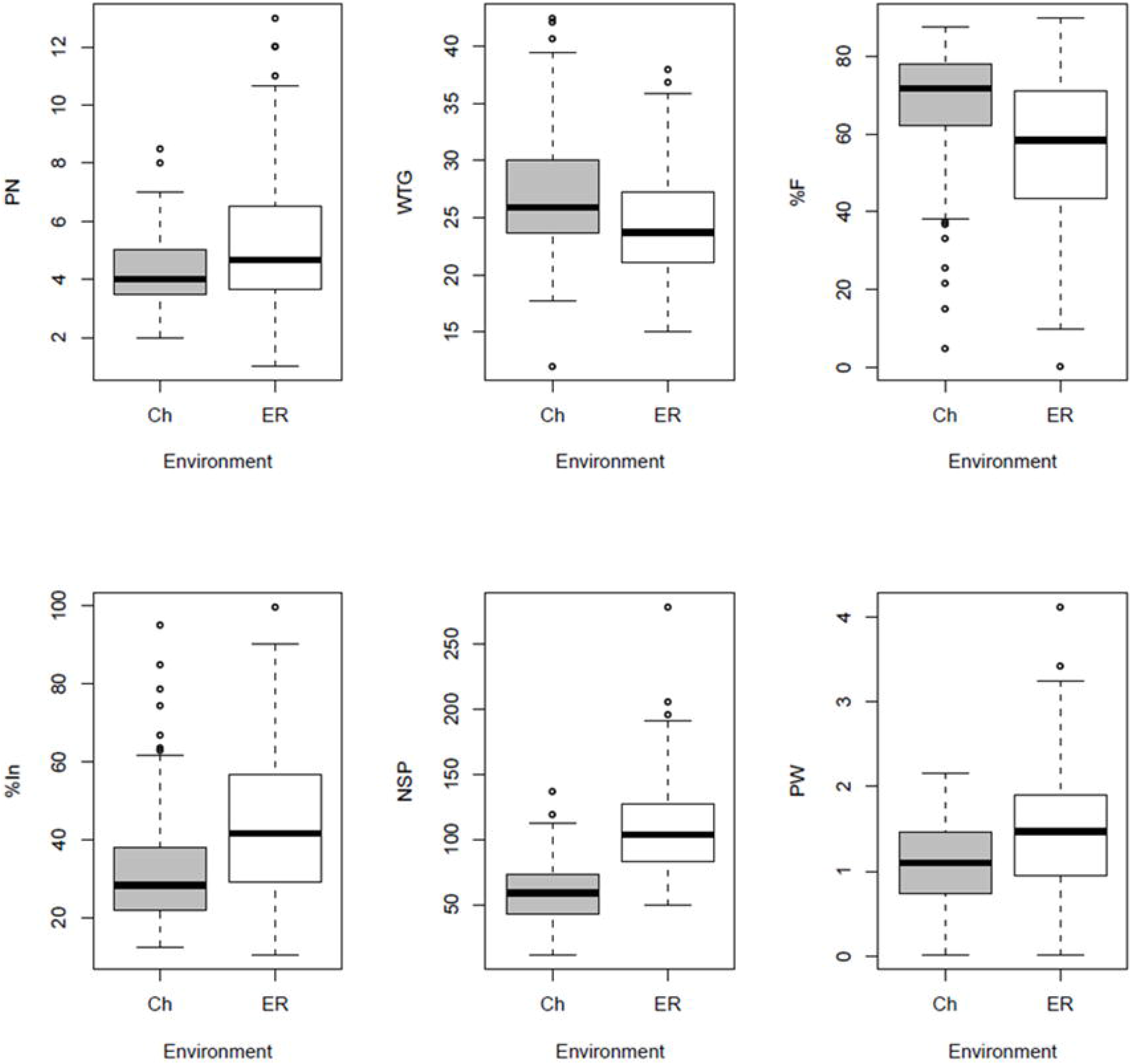
Box diagram of the analyzed yield components. Gray boxes represent plants grown in the CH environment. White boxes represent plants grown in the ER environment. The line inside the box represents the mean value of each phenotype, the top and bottom of each represent the 1st and 3rd quartiles. Open circles represent outliers within the dataset.

The analyses of Pearson’s correlation coefficients were calculated between all the FSII parameters associated with the identified candidate genes (Supplementary Table S3) and panicle traits in both environmental conditions (CH and ER) allowed identifying some significant relationships. The top ranking significantly were: one negative correlation between and a positive correlation with the δ(R_o_) phenotype in CH condition. Sequentially, the δ(R_o_) phenotype also showed a positive correlation with the NSP panicle trait (Supplementary Table S3).

On the other hand, the correlation analyses between, panicle traits and FSII parameters for the ER environment were in general significant among themselves. In line with this, the mean of r Pearsoncoefficient between pairs of fluorescence phenotypes and panicle traits, which were statistically significant was r = 0.14. For this reason, the correlations that presented an r> 0.14 were analyzed (Supplementary Table S3).

From the correlation diagram (Supplementary Table S3), it can be seen that the %In trait correlates positively with the N and DI_o_/RC phenotypes. Also, the NSP trait correlated positively with the REo/RC phenotype, correlated negatively with the PI_total_ phenotype. On the other hand, the PW panicle trait presented 4 significant correlations with fluorescence phenotypes, which is the trait that presented the greatest amount of correlations. The four significant correlations, three were negative with the N, DI_o_/RC, and PI_total_ phenotypes, while the remaining correlation was positive with the REo/RC phenotype.

On the other hand, the WTG and %F traits correlated significantly with the same fluorescence, N, and DI_o_/RC phenotypes; being in both cases negative correlations. Finally, the PN panicle trait only correlated significantly and negatively with the PI_total_ fluorescence phenotype.

## 4. Discussion

Mapping genetic regions that control quantitative features of PSII functionality is a difficult task because photosynthetic processes have a complex and polygenic architecture. Our results showed a clear effect of the environment on the population genetic structure evidenced by the segregation into two groups of different individuals represented by plants grown in the CH and ER environments (Fig. 1).

In plants, chlorophyll fluorescence has been used extensively to examine the influence of abiotic stresses on light-dependent reactions (Fracheboud *et al*., 1999; Huang *et al*., 2004; Silvestre *et al*., 2014; Rachoski *et al*., 2015). However, despite the versatility of the use of chlorophyll fluorescence as a rapid phenotyping method, and of great capacity to evaluate numerous plant accessions, few reports study the mapping of regions or the genetic variation of fluorescence phenotypes in environmental conditions. Nonetheless, QTL for photosynthetic phenotypes associated with light reactions has been successfully mapped in some species of agronomic importance such as wheat, corn, soybeans, and sorghum (Šimić *et al*., 2014; Herritt *et al*., 2016; Ortiz *et al*., 2017).

The reactive processes of the PSII are sensitive to light, and in turn, the light that impacts the PSII depends strongly on the environmental conditions. The natural variations are an important approach to identify new genetic determinants to improve the efficiency of the use of crop light. Since the phenotypic variation of the functionality of the photosynthetic apparatus (PA) of the RDP-1 was measured under normal non-stressful conditions in two environmental locations and the environmental impact on the genetic architecture of the PA could be assessed. Our results showed that fluorescence phenotypes evaluated are highly influenced by genotype, environmental conditions, and genotype-environment interaction (Table 1). Similar results were found in studies in soybean, where chlorophyll fluorescence phenotypes were analyzed in various environments (Herritt *et al*., 2016). On the other hand, in rice, the analysis of other photosynthetic parameters such as stomatal conductance (Gs), net photosynthesis (Asat), and photochemical performance F_v_/F_m_ were affected by environmental, genotypic factors and their interaction between them (Qu *et al*., 2017). In other words, our results show that the significant natural variation in each chlorophyll fluorescence phenotype observed among a plethora of rice genotypes indicates that the selection of these photosynthetic features as an improvement criterion could be possible.

In this sense, of the significant phenotypes found in both environments F_o_/F_m_, F_v_/F_m_, DI_o_/RC, PI_abs_ and δ(R_o_), it was observed that the average values obtained for the ER environment were higher compared to the Ch (Fig. 2A-B). In this line, the F_o_/F_m_ phenotype is the basis of all the expressions derived from the OJIP test. This phenotype allows the phenomenological quantification of the fluorescence behavior of a sample, and at the same time, compares and classifies different types of samples (Strasser *et al*., 2000). The phenotype F_v_/F_m_ is used to quantify the physiological state of plants (Baldassarre *et al*., 2011). The phenotype PI_abs_ indicates the status of FSII, from the absorption of a photon by FSII to the reduction of Q_b_ (Stirbet and Govindjee 2011). Finally, the DI_o_/RC and δ(R_o_) phenotypes explain the energy dissipation by active reaction center and the efficiency of electron transport to the last FSI acceptor respectively (Stirbet *et al*., 2018).

GWAS studies using chlorophyll fluorescence under optimal stress and growth conditions have been reported in soybeans, corn, and sorghum (Hao *et al*., 2012; Strigens *et al*., 2013; Fiedler *et al*., 2016). In all cases, the photosynthetic phenotype features were associated with multiple regions within the genome. Results of the present work yielded 110 QTL in all chromosomes and even the same phenotype was associated in more than one region within the same chromosome according to those previous reports (Fig. 3A; 4A-B). Particularly for the CH environment, a QTL was identified on chromosome 9 for the PI_abs_ phenotype, this phenotype has a unique association on the said chromosome, and in turn, this was the unique association found on this chromosome (Fig. 3B). This fact makes the PI_abs_ phenotype an interesting study target in plant improvement projects since this type of association is an exception to the results conventionally found in GWAS studies for chlorophyll fluorescence phenotypes.

In addition to the difference in the number of SNPs and the phenotypic variation observed between the environments, and as consequence a greater number of QTL for the ER environment were also identified. In line with those mentioned, GWAS studies in Arabidopsis thaliana YELLOW SEEDLING G1 (YS1) revealed diversity in photosynthetic acclimatization under high radiation conditions, finding a greater amount of QŢL than in low radiation conditions (van Rooijen *et al*., 2017).

The analysis of the QTL for the CH environment yielded a single multi-type QTL on chromosome 5 (Fig. 3), unlike the ER environment where 4 multi-type QTLs could be identified (Fig. 4). In both environments, the F_o_/F_m_, F_v_/F_m_, and DI_o_/RC phenotypes share the same QTL on chromosome 5 for plants in CH and chromosome 6 in the ER plants. Identical results were obtained in studies conducted in soybeans (Herritt *et al*., 2016); where the phenotypes of the relationship between maximum and minimum fluorescence, the maximum quantum yield of PSII, and energy dissipation per active reaction center share the same region within the chromosome.

The regions of the genome found by molecular markers for fluorescence phenotypes contained groups of genes that are important for the metabolic and molecular functioning of rice. Additionally, these regions encompassed individual genes that have a significant impact on the functions of grain filling, and therefore, the values of some chlorophyll fluorescence phenotypes could directly influence the grain yield of rice. The examination of the genetic annotations linked to the enrichment analysis of GO terms identified in this study revealed 38 candidate genes (Table 4). Of the identified candidate genes, those with catalytic activity notations formed the majority group (Tables 3 and 4).

When analyzing how fluorescence phenotypes are related to the identified genes, it was observed that the S_m_ and N phenotypes, in general, are associated with the same identified genes (Table 4). Phenotype N indicates the number of reduction and oxidation events of Q_a_ between time zero and Fm (Strasser *et al*., 2000). In turn, the S_m_ phenotype indicates the energy required for electron transport, and thus, closes all reaction centers (Oukarroum and Strasser 2004), this is equivalent to saying that S_m_ indicates the functionality of the quinone complexes (Q_a_/Q_b_). Both phenotypes are highly related to the functioning of the set of quinones, therefore, this may explain that both phenotypes are linked to the same genes. In this sense, of the 5 transcription factors (TF) identified two of them co-located with both phenotypes N and S_m_ (LOC_Os10g39030, LOC_Os05g23780). The functional notation assigned for the first TF corresponded to the homeobox (HD) domain, while a second TF codes for the OsMADS70-MADS-box family of genes with M-alpha type-box (MADS-BOX). Others GWAS reports for panicle traits in RDP-1 populations mapped genes containing both types of HD and MADS-BOX domains associated with flowering time, and the plant height (Begum *et al*., 2015).

The HD domain is a highly conserved motive present in many TFs. At the same time, work carried out with mutant rice plants for HD domains in different TFs showed that those HD domains are essential for the regulation of tillering, sterility and flowering time in rice (Mjomba *et al*., 2016; Wei et al., 2016). On the other hand, in rice, the FT MADS-BOX besides being involved in the specification of the floral organs has also been involved in several aspects of the growth and development of the plants (Arora *et al*., 2007). In this sense, Lee et al. (2004) showed that mutants of the MADS-BOX family of rice presented a delay in flowering, while in rice plants that overexpressed genes of the MADS-BOX family, extremely early flowering was observed.

Of the remaining three TFs, one is associated with the Q_b_RC phenotype (βHelix-loop-helix) and two identified TFs were associated with DI_o_/RC (GATA zinc finger domain and Myb). In all cases, these TFs were associated with different types of abiotic stress. Besides, DI_o_/RC is also linked to TF HD by co-locating with the N and S_m_ phenotypes. Likewise, the DI_o_/RC phenotype presented a large number of identified genes, mainly in plants that were grown in the ER environment. In general, the genes identified were associated only with the DI_o_/RC phenotype. In this sense, the energy dissipation processes have been described as a fundamental mechanism in the photo protection of plants, in response to the excess of light that affects the complex antenna of the chloroplast (Müller *et al*., 2001).

Among the identified candidate genes associated with the DI_o_/RC, the LOC_Os05g44922 and LOC_Os05g44560 genes can be highlighted. For the first of these two genes, the assigned functional notation corresponded to 6-phosphofructokinase (PFK), an intermediate enzyme in the glycolysis pathway conducted by the Calvin cycle. It has been described that PFK is a key intermediary in the synthesis of the main starch component of rice grain (Chang *et al*., 2017). For the second candidate gene, the functional notation corresponded to the genes of the Kinesin family.

In rice, kinesins are linked to cell division processes and are expressed in both pollen grains and somatic tissues. In turn, studies conducted on rice indicated that genes from the kinesin family regulate plant height and seed length (Zhang *et al*., 2010; Kitagawa *et al*., 2010).

Finally, of the two most representative phenotypes evaluated in CH environment, δ(R_o_) was associated with a gene that contained the kinesin motor domain, a similar result was identified for DI_o_/RC phenotype. Furthermore, the PI_abs_ phenotype underlying with LOC_Os09g26620 *loci* that codes for an auxin-repressed protein. Previous reports showed this gene family regulated tillering (Zhang *et al*., 2018). In line with the aforementioned, it has been reported that a large number of tillers limit the remobilization of resources generated by photosynthesis to the final destination, the grain, limiting the yield potential, and increasing biomass (Khush, 1995).

The analysis of panicle traits showed a greater accumulation of positive characters such as PN, NSP, and PW in plants grown in the ER, compared with plants grown in CH (Fig.5). However, even though the plants grown in the ER showed a greater number of destinations in their panicle features, they presented a higher % In (Fig. 5). The higher %In could be that for the ER environment the variability of genotypes that managed to reach the reproductive stage was more diverse within the sub-populations of the RDP-1.

From the correlation diagram for CH (Supplementary Table S3), it was observed that the PI_abs_ phenotype correlated negatively with the NP trait or what is equivalent to saying that when PI_abs_ increase there are fewer panicles per plant. This result coincides with the lower value of NP found in CH plants compared to plants grown in ER (Fig. 5) and with the function described for the identified gene linked to this phenotype. In concordance it has been reported that a greater tendency of plants to increase their tillering allocates photosynthetic resources to generate biomass at the expense of limiting resources destined to the panicle (Khush, 1995). Concerning the δ(R_o_) phenotype, this correlated negatively with NSP and positively with NP. Again, these results are consistent with the underlying gene identified for this phenotype, the kinesins, whose main function is cell proliferation since a higher value of δ(R_o_) has a higher PN, although a lower NSP.

Alternatively inthe ER location, the N parameter evaluated, negatively correlated with the panicle, PW, WTG, and % F characters, positively with the %In trait, in turn, RE_o_/RC positively correlated with NSP and PW (Supplementary Table S3), indicating that a lower number of reduction and oxidation cycles of Q_a_ and greater electron transport favors panicle performance. Finally, the DI_o_/RC phenotype showed the same correlations as for the N phenotype. These phenotypes were linked to genes described as essential for grain yield. Energy dissipation has been described as a photo protection mechanism to optimize photosynthetic efficiency at the canopy level achieving an increase in biomass and yield in rice (Murchie *et al*., 2015).

In conclusion, this work contributed with the identification of genetics determinants of chlorophyll fluorescence diversity in rice by GWAS at two contrasting conditions and how these FSII parameters are related with panicle characteristics. Manifold genes, SNP, and QTL identified here may serve for the design of molecular markers and contribute to the assistance of breeding programs of this species.

The improvement of the photosynthesis capacity in rice constitutes a goal to increased field yields and this work could contribute to it.

## Supplementary data

**Supplementary Figure 1:** Graph of LD vs Genetic distance. The figure shows the graphic analysis of chromosomes 1 to 12.

**Supplementary Figure 2:** Quantile-Quantile graph (QQplot) for the CH environment. The figure shows the phenotype with significant associations.

**Supplementary Figure 3:** Quantile-Quantile graph (QQ plot) for the ER environment. The figure shows the phenotype with significant associations.

**Supplementary Table 1:** Significant associated SNPs markers to PSII identified by GWAS analysis. SNPs linked to their respective fluorescence phenotype, environment and matrix structure are shown.

**Supplementary Table 2:** Total candidate genes identified by GWAS analysis in CH and ER environments.

**Supplementary Table 3:** Correlation diagram between Panicle traits and PSII fluorescence phenotypes for the Ch and ER environment. Asterisks indicate significant correlation according to the Pearson pairwise correlation method (* p <0.05).

## Data availability statement

All data supporting the findings of this study are available within the paper and within its supplementary materials published online.

## Acknowledgments

This work was supported by grant StartUp 2013 0399 provided by Agencia Nacional de Promoción Científica y Técnica (ANPCyT). This work is part of the PhD thesis of JMV.

## Author contribution

JMV, EB, MLP, AAR, and SJM: investigation

JMV, EB, and SJM: formal analysis, writing.

AAR: funding acquisition

AL, JC, and OAR: resources

FC: validation

SJM: conceptualization

## Abbreviations

%In: calculated infertility percentage
%F: fertility percentage
GO: gene ontology
GWAS: genome-wide association
NSP: number of spikelet per panicle
PN: panicle number
PSII: photosystem II
PW: panicle weight
SNP: single nucleotide polymorphisms
WTG: thousand grain weight

